# Social behaviours predict vocal turn-taking in common marmosets

**DOI:** 10.64898/2026.08.11.744109

**Authors:** Yuta Tamai, Jumpei Matsumoto, Julia Löschner, Luca König, Takaaki Kaneko, Ken-ichi Inoue, Koji Toda, Steffen Hage

## Abstract

Human conversation depends on the continuous integration of vocal exchanges with visual and spatial cues, yet the evolutionary origins of this multimodal coordination remain poorly understood. Although vocal turn-taking has been documented across many animal species, studies have largely examined vocal exchanges in isolation from the accompanying social dynamics. Using acoustic localization and 3D pose tracking in freely interacting marmoset pairs, we simultaneously quantified vocal behaviour and social interactions during natural communication. We found that vocal turn-taking is dependent on distinct multimodal behavioural states defined by head orientation, spatial proximity, and ongoing social interaction. While call features did not reliably predict turn-taking, these behavioural dynamics predicted whether vocal exchanges developed into turn-taking or terminated after isolated calls. Our findings reveal that primate vocal communication is fundamentally organized by multimodal behavioural coordination rather than by vocal signals alone, providing an evolutionary framework for understanding the origins of human conversation.

## Introduction

Human social interaction depends on the seamless integration and interpretation of multiple communication cues. During conversation, vocal exchanges are coordinated with eye contact, facial expressions, body orientation, gestures, and changes in interpersonal distance, allowing conversational partners to rapidly adapt to one another ^1–4^. Conversational turn-taking therefore arises from a dynamic interplay between vocal and non-vocal behaviours rather than from acoustic signals alone. Yet, despite its central role in human communication, the evolutionary origins of this multimodal behaviour remain poorly understood.

The common marmoset (*Callithrix jacchus*) provides a powerful model for understanding how behavioural modalities are integrated into vocal turn-taking because they exhibit rich social communication ^5,6^ and flexible vocal behaviour ^7–10^ with conserved features of human vocal communications ^3,7^. Additionally, marmoset vocal behaviour can now be linked to the neural motor circuits that support multimodal communication ^11,12^. However, previous work has predominantly examined antiphonal calling (vocal turn-taking) between spatially separated or visually occluded animals ^7,9,13^. Although these studies established that marmoset vocal exchanges are socially flexible and behaviourally structures, they have largely considered vocal behaviour independently of the accompanying gaze, facial, postural, and spatial cues that may organize it. Consequently, the extent to which these behavioural modalities are coordinated with vocal production during natural, close-range communication among marmosets remains poorly understood.

Here, we combine acoustic source localization with vision-based three-dimensional pose estimation to simultaneously quantify vocal behaviour and three-dimensional social dynamics in freely moving marmosets. Using this integrated approach, these data show that vocal turn-taking is dependent on distinct multimodal behavioural states and that head orientation and spatial proximity shape conversational exchange. Therefore, our findings reveal multimodal behavioural coordination as a fundamental organizing principle of primate communication.

## Results

### Mapping close-range vocal communication in common marmosets

To investigate the types of vocalizations and behaviours marmosets use during antiphonal calling we developed an acoustic camera and multi-view recording system to simultaneously capture vocalizations and the vocalizers between three male-female pairs of common marmosets that are permanently housed together. By ensuring the head of each marmoset and the source of the vocalizations were captured from at least one viewpoint, we reliably identified the vocalizing individual and reconstructed the three-dimensional posture and movements of both animals during natural close-range social interactions (Fig. 1A–C, for detail see Supplementary Methods). We analyzed 1,674 vocalizations and embedded 14 acoustic features (such as call duration, dominant frequency, and frequency range) into a two-dimensional UMAP embedding, revealing seven clusters that grouped into three major call categories: Phee-like (three clusters), Chirp-like (two clusters), and Trill-like (two clusters) (Fig. 1D,E). Trill-like calls were most frequent (1,117 calls, 66.7%), followed by Phee-like calls (351 calls, 21.0%) and Chirp-like calls (189 calls, 11.3%).

**Figure 1.**
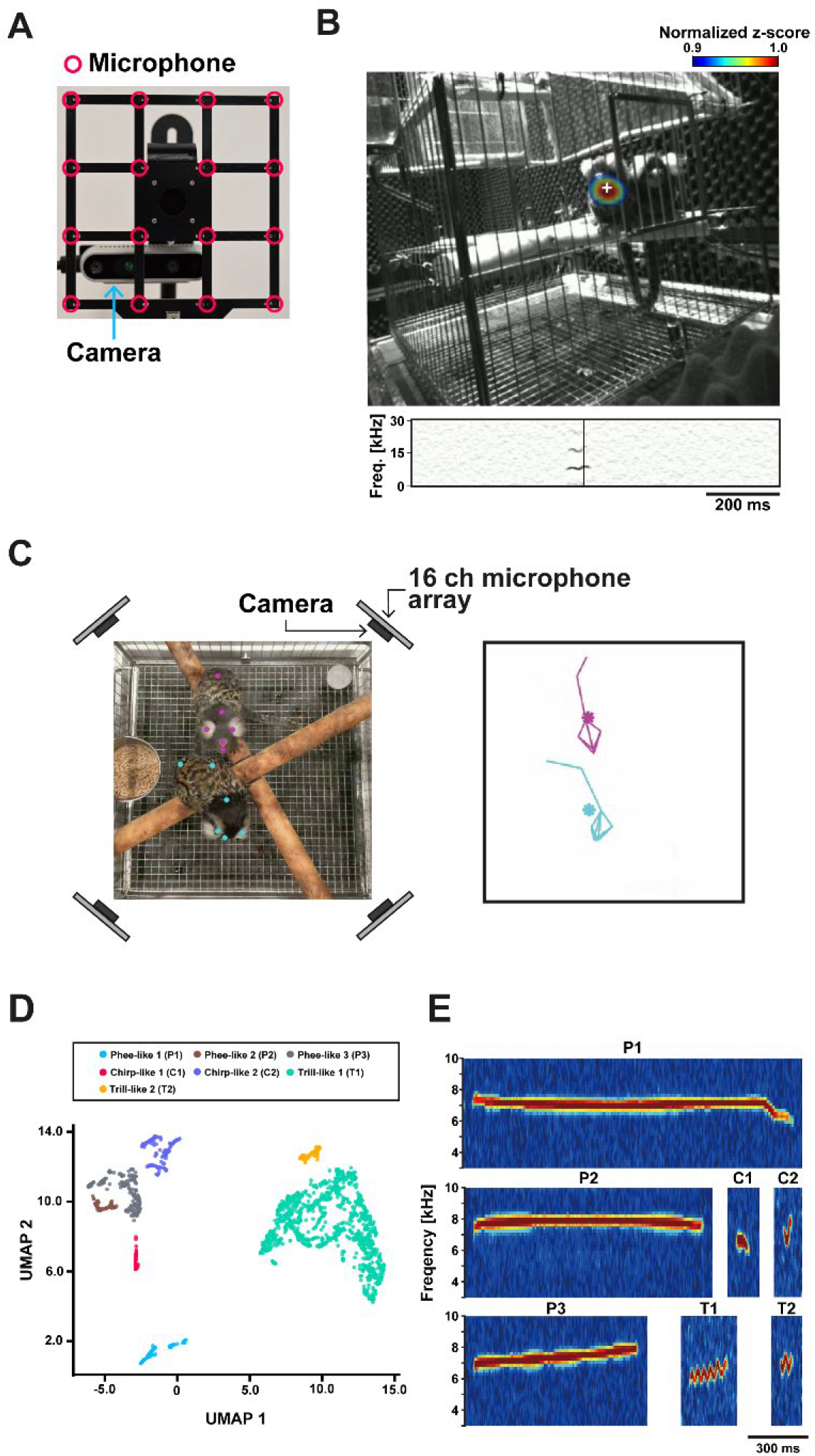
Experimental setup for recording close-range vocal communication in common marmosets. (A) Photograph of the acoustic camera system consisting of a 16-ch microphone array integrated with a camera. The distance of each microphone is 42 mm. (B) Representative example of acoustic source localization during vocal production (top) and the corresponding spectrogram of the detected vocalization(bottom). For visualization, the upper 10% of the frame-wise z-score range was linearly normalized and rendered using the JET colormap. (C) Representative two-dimensional body-part tracking from six synchronized camera views using DeepLabCut (left) and the corresponding three-dimensional reconstruction (right). (D) Two-dimensional UMAP embedding of vocalizations based on 14 acoustic features. Acoustically similar vocalizations were subsequently grouped using HDBSCAN, yielding seven vocalization clusters. (E) Representative spectrograms of the seven vocal clusters identified by HDBSCAN: Phee-like 1, Phee-like 2, Phee-like 3, Chirp-like 1, Chirp-like 2, Trill-like 1, and Trill-like 2.

### Turn-taking interactions are temporally coordinated

Alternating vocal exchanges between paired marmosets were frequent and showed structured turn-taking dominated by Trill-like call sequences (Fig. 2A,B). Trill-like 1 → Trill-like 1 combinations accounted for 60.9% of turn-taking events, followed by Chirp-like 2 → Trill-like 1 sequences (7.6%). Turn-taking latencies, quantified using offset-to-onset (the interval between the offset of the initial call and the onset of the response call) and onset-to-onset (the interval between the onset of the initial call and the onset of the response call) measures peaked at short delays of approximately 300 ms and 400 ms, respectively, and were well-described by a two-component Gaussian mixture model (GMM) that separated short- and long-latency modes (thresholds 797ms and 854 ms; Fig. 2C). Within the short-latency mode, both initial and response-call durations were negatively correlated with offset-to-onset latency (initial call: *r* = −0.260, *P* < 0.01; response call: *r* = −0.250, *P* < 0.01), whereas only response duration correlated with onset-to-onset latency (*r* = −0.298, *P* < 0.001; Fig 2D). Therefore, turn-taking was characterized by rapid and temporally structured vocal responses, with shorter response latencies associated with longer calls, particularly for the responding animal. This relationship suggests that the temporal organization of vocal exchanges depends not only on the interval between calls but also on the duration of the preceding and responding vocalizations.

**Figure 2.**
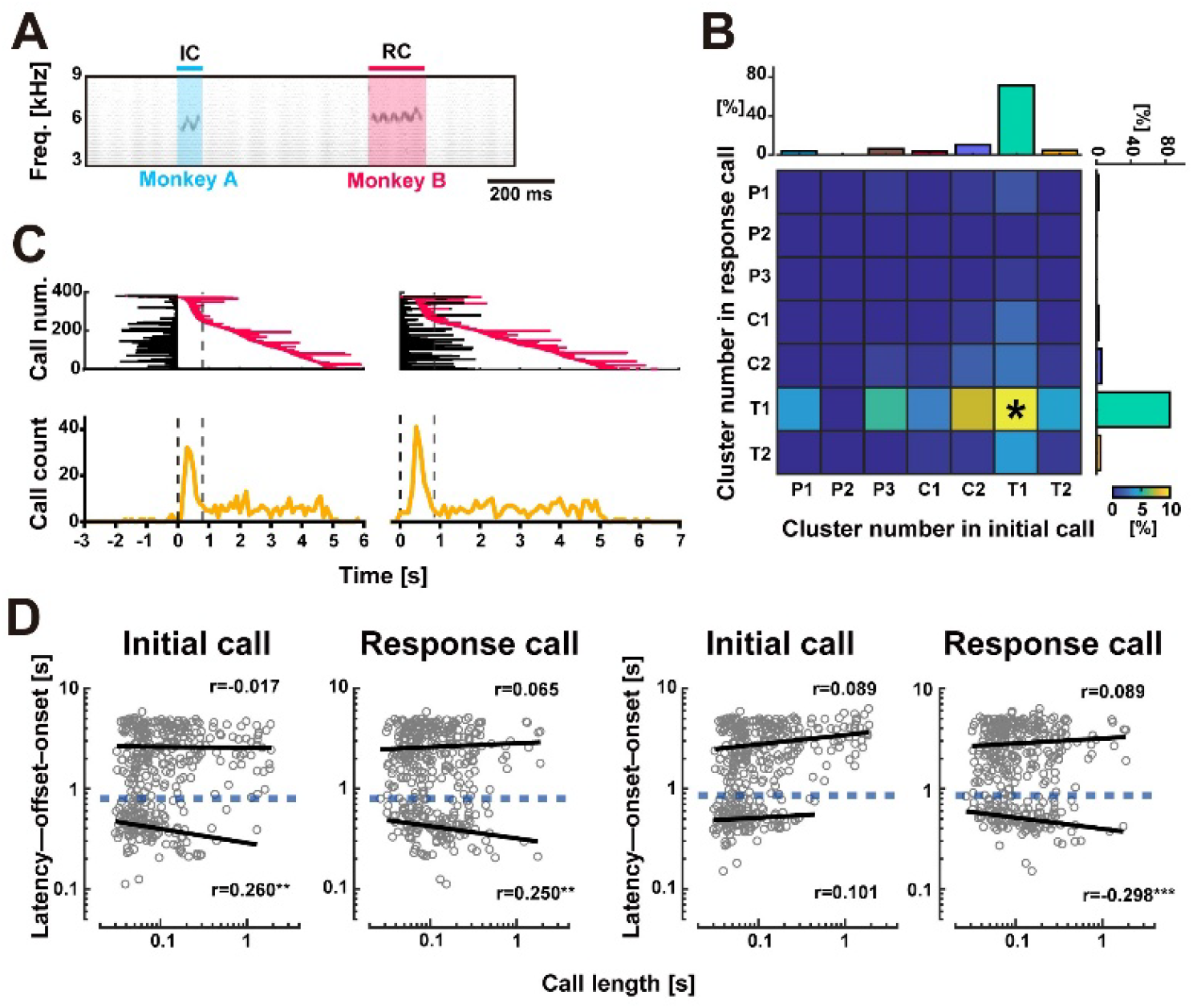
Temporal coordination and characteristics of turn-taking interactions in common marmosets. (A) Representative spectrograms showing vocal exchanges between Monkey A and Monkey B during a turn-taking interaction. (B) Confusion matrix showing combinations of initial and response call types during turn-taking interactions. The asterisk indicates a proportion exceeding the upper limit of the color scale (i.e., 60.9%). (C) Temporal coordination of turn-taking interactions aligned to the offset (left) or onset (right) of the initial call. Grey dashed lines indicate the boundary between short- and long-latency turn-taking interactions identified by a Gaussian mixture model (GMM). (D) Relationship between call duration and response latency (left: initial call, right: response call). Data were divided into short- and long-latency turn-taking interactions using the GMM-derived boundary, and linear regression was performed separately for each group. Response latencies were calculated using the offset-to-onset (left column) and onset-to-onset (right column) intervals. Blue dashed lines indicate the GMM-derived boundary.

### Approach behaviour and head orientation modulates vocal turn-taking

To determine how spatial proximity and ongoing social behaviour shape vocal turn-taking, we examined whether inter-individual distance (IID) trajectories and associated acoustic features distinguished turn-taking (TT) from non-turn-taking (NTT) calls. Social interaction accompanied by vocalizations occurred predominantly at close inter-individual distances (IID), with both TT and NTT calls most frequently produced at 10–25 cm separation (probability> 0.1; Fig. 3A,B). Using DeepLabCut-derived landmark positions, we calculated IID as the moment-by-moment spatial separation between the two animals and extracted IID trajectories from −2s before to 5s after each vocal onset. These IID trajectories were grouped into five clusters using time-series k-means with dynamic time warping (Fig. 3C), and one-way PERMANOVA revealed a significant overall difference in acoustic feature space between TT and NTT calls (*P* < 0.001). Acoustic features of TT calls did not differ across behavioural clusters (*P* > 0.05), whereas NTT calls showed significant cluster-dependent differences (*P* < 0.001). Within-cluster comparisons indicated differences between TT and NTT calls in four of five clusters (Cluster 1: P < 0.01; Cluster 2: P < 0.001; Cluster 3: P < 0.001; Cluster 4: P < 0.05; no difference in Cluster 5; Holm-corrected PERMANOVA). Consistent with these multivariate differences, acoustic features (call duration, dominant frequency, and inflection point count) were stable in TT calls and more variable in NTT calls across clusters, suggesting that vocal features are relatively constrained during reciprocal exchanges, whereas their expression is more strongly modulated by the surrounding behavioural and spatial context when calls do not elicit a vocal response. Thus, acoustic structure alone may be insufficient to determine whether a call participates in turn-taking; rather, call features appear to acquire their communicative function within an ongoing social interaction. Because Cluster 5 was characterized by a progressive decrease in IID following vocal onset, we next asked which animal initiated the approach and found that the initial call vocalizer was more likely to approach their partner, whereas responding callers were more likely to remain stationary while being approached by the initial caller during turn-taking interactions. This asymmetry suggests that vocal turn-taking is accompanied not only by coordinated vocal timing but also by role-dependent coordination of approach behaviour (Fig. 3E).

**Figure 3.**
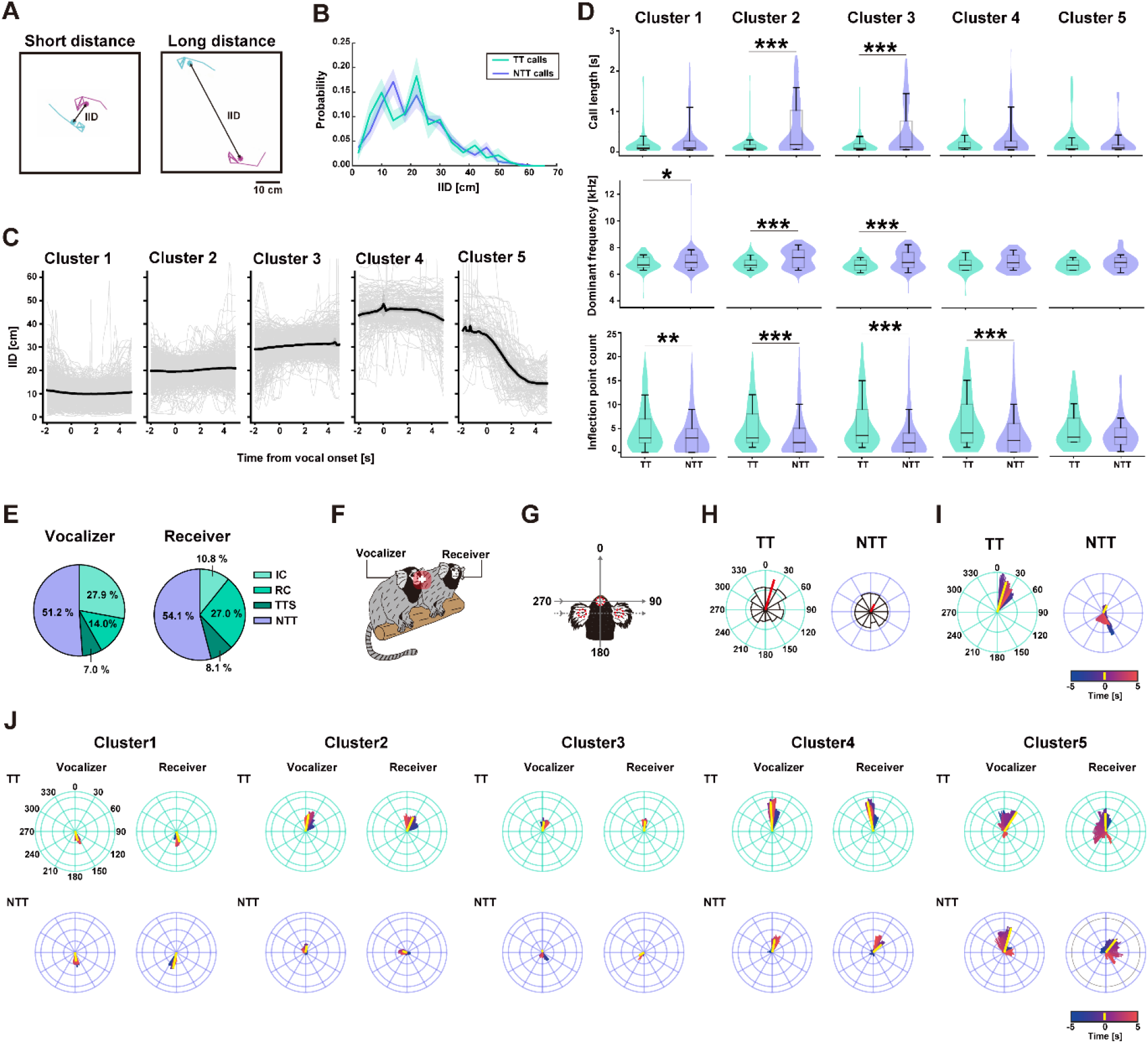
Inter-individual distance and Head-direction dynamics during turn-taking vocal interactions in common marmosets. (A) Representative three-dimensional reconstructions illustrating short and long inter-individual distances (IID) between two marmosets. (B) Normalized histograms of IID during turn-taking (TT) and non-turn-taking (NTT) vocalizations. Solid lines represent the mean normalized probability across animals, and shaded areas indicate the SEM. Time courses of IID from 2 s before to 5 s after vocal onset. IID trajectories were clustered into five behavioural clusters based on Dynamic Time Warping (DTW) distances. Black and gray lines indicate the cluster means and individual trajectories, respectively. (D) Comparison of acoustic features (call duration, dominant frequency, and inflection point count) between TT and NTT vocalizations within each behavioural cluster. *, *P*< 0.05; **, *P*< 0.01; ***, *P*< 0.001; two-tailed Mann–Whitney U test with Holm correction for multiple comparisons. Violin plots show the data distribution. Box plots show the median (center line), interquartile range (box), and the 10th–90th percentiles (whiskers). (E) Pie charts showing the proportions of communication types underlying the approach behaviour observed in Cluster 5, in which IID decreased following vocal onset. Approach events were categorized according to whether the caller or the receiver approached the interaction partner. The pie charts indicate the proportions of Initial call (IC), Response call (RC), Turn-taking sequence (TTS), and Non-turn-taking call (NTT) within each approach category. (F) Schematic illustrating the definitions of the vocalizer and receiver during vocal interactions. (G) Definition of head direction. The head orientation vector was defined from the midpoint between the left and right ears to the forehead. Head direction was quantified as the angle between the head orientation vector and the vector from the forehead to the centroid of the interaction partner, measured in the plane defined by the forehead, left ear and right ear. Red dashed circles indicate the forehead, left ear and right ear landmarks. The dashed line connecting the ears represents the reference plane. (H) Polar plots of the vocalizer’s head direction at vocal onset during turn-taking (TT) and non-turn-taking (NTT) vocalizations. Red vectors indicate the mean resultant vectors, with their directions representing the circular means and their lengths representing the mean resultant lengths. Concentric circles indicate normalized vector lengths in increments of 0.1. (I) Time-resolved mean resultant vectors of the vocalizer’s head direction from 5 s before to 5 s after vocal onset during TT (left) and NTT (right) vocalizations. Colors indicate time relative to vocal onset. (J) Time-resolved mean resultant vectors of the head directions of the vocalizer and receiver for each behavioural cluster shown in (C). Colors indicate time relative to vocal onset.

Vocal turn-taking was accompanied by characteristic head-direction patterns of both the vocalizing animal (vocalizer) and its partner (receiver) (Fig. 3F, G). During TT calls, vocalizer head direction was significantly oriented toward the partner animal (Rayleigh test, P < 0.001), whereas no directional bias was observed during NTT calls (Rayleigh test, P = 0.134; Fig. 3H). Angular head-direction distributions differed significantly between TT and NTT vocalizations (Mardia–Watson– Wheeler test, P < 0.01; Fig. 3H), and vocalizers consistently oriented toward their partners around TT calls but not NTT calls (Fig. 3I). Across behavioural IID clusters, Cluster 1 (short IID) showed mutual head orientation away from each other; Clusters 2 and 3 (intermediate IID) showed mutual orientation in the same direction during TT but not NTT, and Clusters 4 and 5 (larger or decreasing IID) showed general orientation toward each other regardless of interaction outcome (Fig. 3J). Therefore, head orientation was coupled to vocal turn-taking in a spatially dependent manner: partner-directed alignment characterized TT calls at intermediate distances, whereas orientation patterns at very short and larger distances were more strongly determined by spatial configuration than by vocal exchange outcome.

### Head orientation and proximity predict turn-taking vocalizations

To ascertain the importance of social behavioural interactions in determining antiphonal calling, we next asked if they predict whether vocal interactions develop into TT or NTT. Logistic regression based on IID and head-direction features revealed two prominent decoding peaks at approximately 2.5 s before vocal onset (P1) and 2.0 s after vocal onset (P2), with an additional increase at vocal onset (VO; Fig. 4A). At these time points confusion matrices confirmed reliable classification, whereas performance in control windows (±5 s) was near chance (Fig. 4B) and balanced-accuracy decoding performance exceeded chance based on permutation testing (P1: P < 0.01; VO: P < 0.05; P2: P < 0.001; Fig. 4 C, D), indicating temporally localized predictive information. In contrast, decoding based on acoustic features alone did not significantly exceed chance performance (BA = 0.524, P = 0.131). Feature selection based on the frequency of non-zero coefficients from L1-regularized logistic regression across leave-one-subject-out cross-validation folds showed that vocalizer head-orientation features were consistently selected for TT classification at P1, VO, and P2. Receiver head-orientation features contributed primarily around vocal onset, whereas IID-related features emerged at P2 (Fig. 4E), indicating that head-orientation is the most robust predictor, with spatial proximity contributing after vocalization onset.

**Figure 4.**
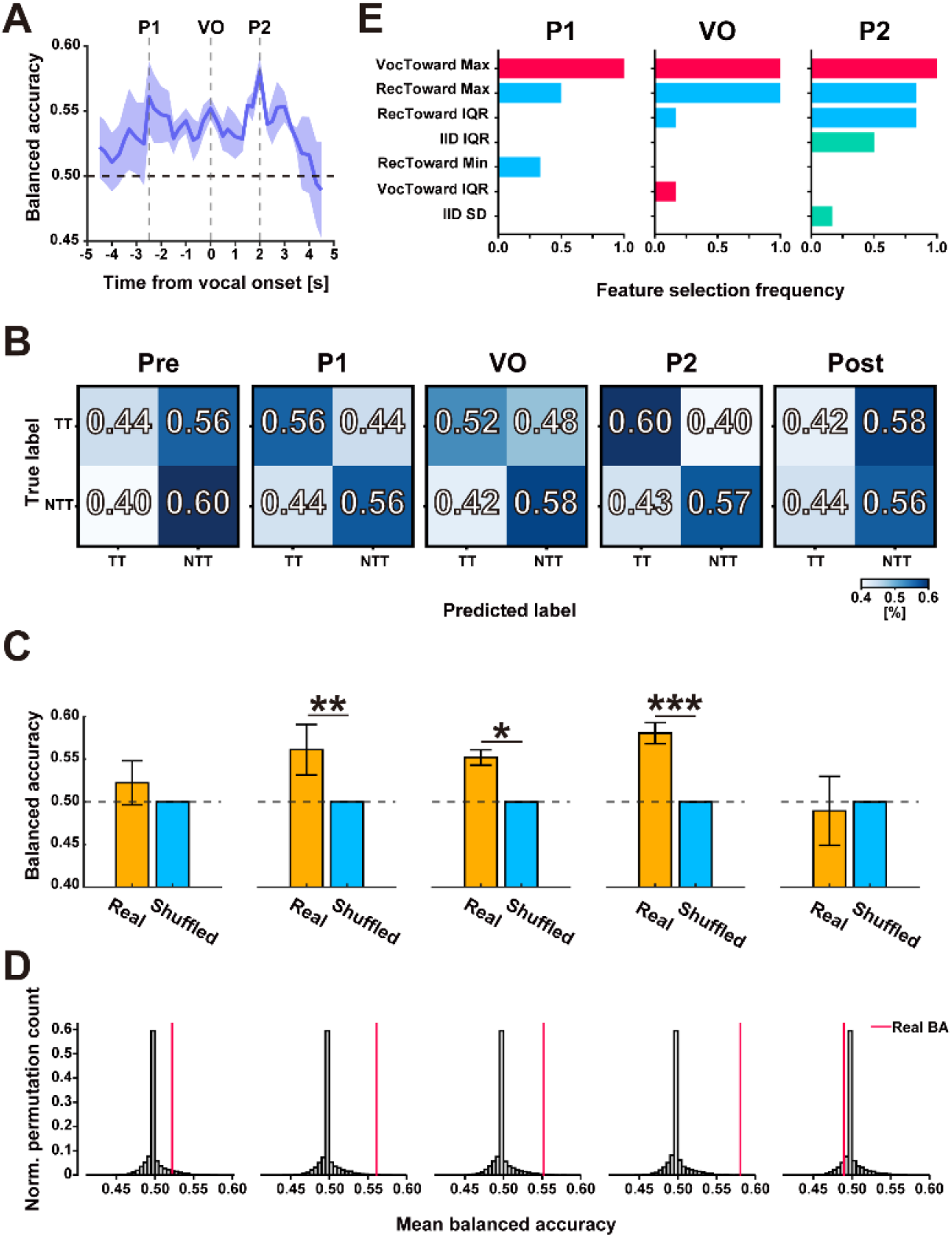
Behavioural decoding of turn-taking from inter-individual distance and head-orientation features. (A) Time course of decoding performance for predicting whether a vocal interaction developed into turn-taking (TT) or non-turn-taking (NTT) using behavioural features extracted from sliding temporal windows. The solid line indicates mean balanced accuracy (BA) across subjects, and the shaded area represents SEM. Dashed vertical lines indicate the three representative temporal windows: peak 1 (P1, −3 to −2 s), vocal onset (VO, −0.5 to 0.5 s), and peak 2 (P2, 1.5 to 2.5 s). The black horizontal dashed line indicates chance-level performance (BA = 0.5). (B) Subject-normalized confusion matrices for the corresponding temporal windows (Pre, P1, VO, P2, Post). Pre and Post denote the temporal windows −5 to −4 and 4 to 5, respectively. Diagonal values indicate correct classification rates, whereas off-diagonal values indicate misclassification rates. (C) Mean BA for the representative temporal windows (Pre, P1, VO, P2, and Post) obtained using the original labels (Real) and label-shuffled data (Shuffled). Error bars indicate SEM across subjects. Gray dashed lines indicate the chance level (BA = 0.5). *: *P*< 0.05; **: *P*< 0.01; ***: *P*< 0.001 (permutation test with Holm correction for multiple comparisons). (D) Null distributions of BA obtained from 10,000 label permutations for each representative temporal window (Pre, P1, VO, P2, and Post). Red vertical lines indicate the observed decoding performance obtained from the original data. (E) Feature selection frequency with which behavioural features were selected by the L1-regularized classifier across leave-one-subject-out cross-validation (LOSO-CV) folds in the three representative temporal windows (P1, VO, and P2). Only features selected in at least one-fold are shown. Colors indicate feature categories: red, vocalizer-toward features; blue, receiver-toward features; green, inter-individual distance (IID) features.

## Discussion

Here, we demonstrate that vocal turn-taking in marmosets is not a vocal phenomenon alone. Instead, antiphonal calling emerges from multimodal behavioural states shaped by head-orientation and spatial proximity. Although acoustic features contributed to distinguishing TT from NTT calls, they did not provide fixed markers of turn-taking: they were stable across behavioural contexts during TT calls but more variable during NTT calls. This indicates that vocal structure is shaped by the surrounding spatial and behavioural context and functions together with, rather than independently of, non-vocal behaviour. By combining acoustic source localization with three-dimensional pose tracking, we show that reciprocal vocal exchanges are organized by coordinated behaviour rather than by vocal timing in isolation. This finding provides direct experimental evidence for a framework of multimodal, embodied coordination in close-range social communication.

Earlier work showed that marmosets exchange long-distance calls in structured sequences and that turn-taking is influenced by learning, individual calling rate, and social context such as sex and status ^7,13–16^. Together, these studies established that turn-taking is socially flexible and behaviourally structured, and they pointed to the possibility that vocal exchange is embedded in a broader interactional context. However, that multimodal framework remained largely inferential, mainly because most research so far focused on long-distance communication between animals interacting acoustically while out of sight ^7,13,14^. Our data now provide direct evidence that close-distance vocal behaviour becomes conversational based on features of social interactions. At the distances where interactions were most frequent (10–25 cm), turn-taking calls showed relatively stable acoustic features across behavioural contexts, whereas non-turn-taking calls varied with the surrounding spatial and behavioural state. Turn-taking was also accompanied by role-dependent approach behaviour, with initiating callers more likely to approach and responding callers more likely to remain stationary. Thus, whether a call participates in an exchange cannot be determined from its acoustic structure alone; its function emerges from the temporal, spatial, and behavioural context in which it is produced. In this sense, the relevant unit of analysis is not the isolated call, but the social interaction surrounding it.

Head-orientation, rather than acoustic features of calls, emerged as the most consistent predictor of whether vocal interaction developed into turn-taking. Accordingly, decoding based on acoustic features alone did not significantly exceed chance performance. When callers were not in physical contact, vocalizers oriented toward their partners when calls led to exchange, whereas non-turn-taking calls lacked this directional coupling. This suggests that turn-taking is prepared through social alignment before vocal onset. Although call features were relatively stable in turn-taking calls and more variable in non-turn-taking calls, these acoustic differences alone did not reliably distinguish whether vocal interactions developed into turn-taking. Vocal signals therefore acquire their function within a behavioural framework that precedes and shapes their use. Behavioural decoding analyses further support this interpretation. Significantly predictive behavioural information emerged before vocal onset, peaked around the onset of the call, and remained above chance afterward, indicating that head-orientation consistently contributed to distinguishing TT from NTT interactions, whereas spatial proximity contributed after vocal production had begun. Early orientation likely reflects communicative readiness, whereas later spatial dynamics may determine whether exchange continues or ends. Taken together, these results point to a continuous coupling of movement, orientation, and vocal output, rather than a simple stimulus-response sequence.

Understanding how this multimodal coordination is implemented in the brain represents an important next step. Recent ultra-high-field fMRI work has identified a distributed marmoset brain network that integrates vocal and facial social signals, involving auditory and face-processing regions as well as parietal, prefrontal, and cingulate areas ^17–19^. By linking these neural systems to naturally occurring vocal exchanges and the accompanying orientation and spatial dynamics, our findings establish the marmoset as a particularly tractable model for investigating how neural circuits support multimodal social communication and vocal turn-taking. These findings place marmoset communication in a broader comparative framework for understanding how coordinated social behaviour can shape vocal exchange. Although human conversation is more complex and depends on additional linguistic and social mechanisms ^20,21^, our results show that close-range vocal turn-taking in marmosets is organized by multimodal cues, including head-orientation and spatial context. This supports the idea that the conversational dynamics observed in marmoset vocal interactions may arise from embodied social coordination, potentially representing an evolutionary precursor to—but not being directly equivalent to—human conversation. ^20^.

More broadly, our results argue that comparative studies of communication must move beyond vocalizations in isolation. Rather, call function emerges from the surrounding social and spatial context, consistent with broader accounts of communication as a multimodal, interactional system ^22,23^. By linking close-range vocal exchange to dynamic social behaviour, our findings provide a framework for studying the origins and evolutionary precursors of conversation-like behaviour and its neural basis.

## Acknowledgments

This study was supported by DFG Research Unit 5768: HA 5400/6-1 – 532521431 (to S.R.H.), DFG Grant LO3184/1-1 (to J.L.) the Japan Society for the Promotion of Science (JSPS) KAKENHI Grants Nos. 24KK0210 (to Y.T.), 24KJ1927 (to Y.T.), 26K23858 (to Y.T.), 22H05157 (to K.I. and J.M.), Japan Science and Technology Agency (JST) FOREST Program JPMJFR2320 (to J.M.), and Keio Academic Development Fund (to K.T.).

## Author contributions

Conceptualization: Y.T., and S.R.H.; Methodology: all authors; Investigation: Y.T., and J.L.; Visualization: Y.T.; Funding acquisition: Y.T., K.T., and S.R.H.; Project administration: S.R.H.; Supervision: S.R.H.; Writing – original draft: Y.T., and S.R.H. Writing – review & editing: all authors.

## Data availability

All data needed to evaluate the conclusions in the paper are present in the paper. Additional data related to this paper may be requested from the corresponding author.

## Methods

### Experimental animals

Three adult male-female pairs of common marmosets (Callithrix jacchus), each permanently housed together, were used in this study. All animals were born in captivity and maintained as stable pairs throughout the experiments. At the start of the experiments, the animals were aged 8-9 years. Animals had ad libitum access to water and were maintained on a restricted feeding schedule consisting of commercial pellets, supplemented daily with fruits, vegetables, mealworms, and locusts. Additional treats (e.g., marshmallows) were used as positive reinforcement to facilitate transfer from the home cage to the experimental cage. Environmental conditions were maintained at 26°C, 40-60% relative humidity, and a 12 h:12 h light/dark cycle, including simulated dawn and dusk transitions. All procedures were conducted in accordance with institutional and governmental guidelines for animal experimentation and were approved by the Regierungspräsidium Tübingen.

### Acoustic camera and sound source localization

To identify the vocalizing individual during natural vocal interactions, we developed an acoustic camera system for marmoset monkeys based on a sound source localization framework described earlier for mice ^24^. In contrast to the original system, which used a different microphone configuration, our setup employed four sound camera units positioned around the sides of the behavioural arena, each consisting of a 16-channel microphone array (UMA-16 v2, miniDSP Ltd., Hong Kong, China) and a synchronized video camera (Intel RealSense Depth Camera D435, Intel Corporation, Santa Clara, CA, USA). This arrangement ensured that the head and mouth region of each freely moving marmoset were captured from at least one viewpoint, enabling reliable assignment of localized vocalizations to the emitting individual. The sound source localization algorithm followed the previously described approach: time differences of sound arrival across microphones were used to estimate the spatial likelihood of the sound source with a conventional Bartlett (delay-and-sum) beamformer ^24,25^. The resulting sound localization map was synchronized with the video recording, allowing the estimated sound source position to be overlaid onto behavioural images. Vocalizations were assigned to the individual whose head position was closest to the estimated sound source at the time of vocal production. Localization results were subsequently validated by visual inspection.

### Experimental design and setup

During each recording session, animals freely interacted within the experimental cage while synchronized audio and video data were acquired. Behaviour was simultaneously recorded from six viewpoints. Two cameras (Logitech Brio 4K) controlled by the Tucker-Davis Technologies (TDT) system were positioned above and obliquely above the cage to capture overhead perspectives. Vocalizations were recorded using a microphone (MKH 8020 microphone with MZX 8000 preamplifier, Sennheiser, Germany) and digitized using an A/D interface (RX8, Tucker-David Technologies, U.S.A.). In addition, four acoustic camera units were positioned around the sides of the cage, providing lateral views from four directions. Each acoustic camera system simultaneously acquired synchronized audio and video data and operated independently of the TDT system. This multi-view recording setup enabled concurrent monitoring of vocalizations and social behaviours from complementary perspectives throughout each session.

### Vocal call detection and acoustic feature extraction

Vocalizations were automatically detected using USVSEG ^26^, following previously described procedures. Continuous audio recordings were converted into spectrograms, and vocalization segments were identified based on spectral peak continuity after background noise suppression. The detection frequency range was adjusted to preferentially capture marmoset vocalizations with first harmonics between 4 and 16 kHz, based on previous studies, including own (Agamaite *et al*., 2015; Gultekin *et al*., 2021). All automatically detected segments were visually inspected, and false-positive detections were manually removed before subsequent analyses.

Following segmentation, acoustic features were quantified using custom-written MATLAB scripts. For each vocalization, 14 acoustic features were extracted: call duration, dominant frequency, minimum and maximum frequencies, frequency range (max-min frequency), temporal positions of the minimum and maximum frequencies within the call, start and end frequencies, frequency slope, Wiener entropy, number of frequency inflection points, modulation depth, and modulation frequency. Spectrograms were computed using a 512-point Fast Fourier Transform (FFT) with a Hanning window. The extracted features were used for subsequent analyses of vocalization characteristics, whereas detected vocalization onset and offset times were used for sound source localization and synchronization with the behavioural video recordings, enabling identification of the vocalizing individual using the acoustic camera system described below.

### Acoustic feature embedding and clustering

To visualize the distribution of vocalizations in acoustic feature space and identify groups of acoustically similar vocalizations, acoustic features extracted from each vocalization were subjected to dimensionality reduction and clustering. For vocalizations with zero or one inflection point, modulation depth and modulation frequency were undefined and were therefore set to zero prior to analysis. The 14 acoustic features were then standardized using z-score normalization prior to dimensionality reduction. Uniform Manifold Approximation and Projection (UMAP) was then applied to generate a two-dimensional embedding while preserving the structure of the feature space. UMAP was implemented using default parameters with a fixed random seed. The resulting embeddings were subsequently clustered using Hierarchical Density-Based Spatial Clustering of Applications with Noise (HDBSCAN). Clustering was performed with min_cluster_size = 50, min_samples = 10, and the Euclidean distance metric, allowing the algorithm to automatically identify clusters of acoustically similar vocalizations while classifying isolated vocalizations as noise. The resulting cluster assignments were used for subsequent analyses of vocalization categories. Only six calls (0.4 %) were classified as noise and were excluded from subsequent analyses.

### Turn-taking identification and vocal interaction analysi

Turn-taking vocal interactions were identified based on the temporal structure of the calls produced by the two animals. A vocal interaction was classified as a turn-taking event when one individual produced vocalization within 5s of call from another individual. Within each turn-taking event, the first call was defined as the initial call and the subsequent call as response call. Two latency measures were calculated to characterize turn-taking behaviour: (1) Latency (offset-onset), defined as the interval between the offset of the initial call and the onset of the response call, and (2) Latency (onset-onset), defined as the interval between the onset of the initial call and the onset of the response call. To characterize distinct temporal patterns of vocal responses, two-component Gaussian mixture models (GMM) were fitted separately to the distributions of non-negative offset-to-onset and onset-to-onset latencies. For each latency measure, the two Gaussian components were designated as short- and long-latency components according to their estimated means. The boundary between the two components was defined as the latency at which their posterior probabilities were equal, and each turn-taking event was classified as a short- or long-latency interaction according to this boundary. To examine vocal exchange patterns, the acoustic call types of initial and response calls were assigned based on the HDBSCAN clustering of the UMAP embedding described above. Combinations of call types exchanged during turn-taking interactions were then quantified to assess the structure of vocal communication between individuals.

Vocalizations not followed by a response from the other individual within 5 s were classified as non-turn-taking (NTT) vocalizations and analyzed separately.

### Behavioural recording and feature extraction

Behavioural interactions were recorded using multiple synchronized camera systems as described above. Recorded videos were analyzed using DeepLabCut ^27^, a deep learning-based, vision-based pose estimation toolbox. Seven anatomical landmarks were tracked for each animal: forehead, left ear, right ear, nose, occipital region, back, and hip. These tracked landmarks were used to quantify spatial positions and head orientations of both animals throughout the recording sessions. Derived from these landmarks, behavioural variables describing social interactions – including inter-individual distance (IID) and the relative orientation between caller and receiver – were computed and used for subsequent behavioural analyses and machine-learning-based decoding.

Three-dimensional coordinates of the tracked anatomical landmarks for both marmosets were reconstructed by integrating vision-based pose estimates from the six synchronized camera views using “Anipose” ^28^, following the reconstruction pipeline described by our previous study ^29^. Based on the tracked anatomical landmarks, the centroid of each animal was computed for every video frame. Inter-individual distance (IID) was defined as the Euclidean distance between the centroids of the two animals and was calculated throughout each recording session. For each animal, the head direction vector was defined as the three-dimensional vector from the midpoint between the left and right ears to the forehead. Orientation relative to the interaction partner was quantified as the angle between this head direction vector and the vector extending from the animal’s forehead to the centroid of the other animal. Head direction was computed separately for the vocalizing animal (caller) and the responding animal (receiver). To characterize behavioural dynamics associated with vocal communication, temporal changes in IID, caller head direction, and receiver head direction were extracted from 5 s before to 5 s after vocalization onset. These time series were used to compute summary behavioural features for each analysis window and served as input variables for the machine-learning analyses described below.

### Behavioural trajectory clustering

Behavioural responses were further characterized by clustering the temporal dynamics of inter-individual distance (IID). For each vocalization, the IID trajectory from 2 s before to 5 s after vocal onset was extracted. Pairwise similarities between IID trajectories were quantified using Dynamic Time Warping (DTW), allowing comparison of time-series with temporal variability. To accommodate modest variability in the timing of behavioural responses while avoiding excessive temporal warping, DTW alignment was constrained using a Sakoe– Chiba band with a radius of five samples (0.5 s), allowing temporal shifts within ±0.5 s. IID trajectories were then clustered using time-series k-means with DTW as the distance metric. To determine an appropriate number of clusters, candidate solutions were evaluated according to inertia, silhouette scores computed from the pairwise DTW distance matrix, cluster-size distributions, and the interpretability of the resulting behavioural patterns. Based on the combined evaluation assessment of these criteria, a five-cluster solution was selected for subsequent analyses. The resulting clusters were used for subsequent visualization and comparison of vocal interaction patterns.

### Behavioural decoding analysis

To determine whether social vocal interactions would develop into turn-taking (TT) or non-turn-taking (NTT), we performed binary classification using L1-regularized logistic regression. For each vocalization, behavioural features extracted from sliding temporal windows served as predictors, and vocal interaction outcome (TT or NTT) as the binary response variable. The feature set comprised seven IID descriptors (mean, median, standard deviation, slope, 10th percentile, 90th percentile, interquartile range) and fourteen head-orientation (Toward) descriptors. For decoding analyses, “Toward” was defined as the cosine of the angle between the head-orientation vector and the vector pointing to the interaction partner, yielding values ranging from −1 to 1, with values approaching 1 indicating that the animal was oriented more directly toward its interaction partner. For both caller and receiver, seven summary descriptors (mean, median, standard deviation, slope, minimum, maximum, interquartile range) were calculated, yielding 21 features in total. Decoding was performed over a time window from −5 to +5 s relative to vocal onset using a sliding-window approach. A 1-s window was shifted in 250-ms increments (37 overlapping windows), and features were computed independently for each window. A separate classifier was trained for each window. Logistic regression was implemented using the scikit-learn library with L1 regularization and the SAGA optimizer. Features were standardized using z-score normalization (StandardScaler), and class imbalance was addressed using balanced class weights. The regularization parameter was fixed at C = 0.0223 based on a two-stage grid search. Candidate C values were first evaluated over a broad logarithmic range and then refined within a narrower range around the best-performing region using leave-one-subject-out cross-validation (LOSO-CV). For each candidate C value, mean balanced accuracy was calculated across all temporal windows, and the value yielding the highest overall mean balanced accuracy was selected and applied uniformly to all decoding analyses. Generalization performance was evaluated using leave-one-subject-out cross-validation (LOSO-CV). In each fold, all vocalizations from one subject were used for testing and those from the remaining subjects for training, iterating until each subject served as the test set once. Statistical significance of decoding performance was assessed using permutation tests for representative temporal windows. Class labels (TT/NTT) were randomly permuted while preserving behavioural features, and the decoding procedure was repeated 10,000 times. For each window, the mean balanced accuracy (BA) across subjects was computed to form a null distribution. Observed performance was compared against this distribution using a one-sided test, with P values corrected for multiple comparisons using the Holm method. Classifier performance was further characterized using confusion matrices derived from LOSO-CV predictions. For each subject, predicted and true labels were used to compute a confusion matrix, which was normalized across true classes (row normalization) to ensure equal weighting across subjects. Normalized matrices were then averaged to obtain a mean subject-normalized confusion matrix, where diagonal elements indicate correct classification rates and off-diagonal elements indicate misclassification rates. To identify features contributing to TT/NTT decoding, feature selection was examined in three representative temporal windows showing increased decoding performance (P1: −3 to −2 s, VO: −0.5 to 0.5 s, and P2: 1.5 to 2.5 s relative to vocal onset). Regression coefficients were extracted from each LOSO-CV training fold. Given that L1 regularization induces sparsity, features with absolute coefficients greater than 1 × 10^−8^ were considered selected. Selection frequency was defined as the proportion of folds in which a feature was selected. Features selected in at least one-fold across any of the three windows were visualized. Selection frequency was used as an index of feature-selection stability across LOSO-CV folds, reflecting how consistently each feature contributed to the behavioural decoding.

### Acoustic decoding analysis

To determine whether acoustic properties of vocalizations distinguish TT from NTT interactions, we performed binary classification using the same L1-regularized logistic regression framework as that used for behavioural decoding. The 14 acoustic features described above served as predictors. For vocalizations containing zero or one inflection point, modulation depth and modulation frequency were undefined and were therefore set to zero prior to decoding. The regularization parameter was determined using the same two-stage grid-search procedure, yielding C = 1.5199. Decoding performance and statistical significance were evaluated using the same LOSO-CV and permutation procedures (10,000 permutations) as described for the behavioural decoding analysis.

### Statistical analysis

Statistical analyses were performed using custom-written scripts in Python (version 3.8.10) and MATLAB (version R2021b). Statistical tests used for individual analyses are specified in the corresponding Methods sections and figure legends. Unless otherwise stated, all tests were two-sided. Decoding performance was assessed using one-sided permutation tests with 10,000 label permutations. Where applicable, P values were adjusted for multiple comparisons using the Holm method. Statistical significance was defined as P < 0.05.

